# Ontogeny of learning and visual discrimination in zebrafish

**DOI:** 10.1101/2022.02.09.479666

**Authors:** Maria Santacà, Marco Dadda, Luisa Dalla Valle, Camilla Fontana, Gabriela Gjinaj, Angelo Bisazza

**Affiliations:** Department of Biology, University of Padova, Padova, Italy; Department of General Psychology, University of Padova, Padova, Italy; Padua Neuroscience Center, University of Padova, Padova, Italy

## Abstract

With the exception of humans, early cognitive development has been thoroughly investigated only in precocial species, well developed at birth and with a broad behavioural and cognitive repertoire. We investigated another highly altricial species, the zebrafish, *Danio rerio*, whose embryonic development is very rapid: 72 hours. The nervous system of hatchlings is poorly developed, and their cognitive capacities are largely unknown. Larvae trained at 8 days post-fertilisation rapidly learned to associate a visual pattern with a food reward, showing significant performance at 10 days post-fertilisation. We exploited this capacity to study hatchlings’ discrimination learning capacities. Larval zebrafish rapidly and accurately learned colour and shape discriminations. They also discriminated a figure from its mirror image and from its 90°-rotated version, although with lower performance. Our study revealed impressive similarities in learning and visual discrimination capacities between newborn and adult zebrafish despite their enormous differences in brain size and degree of development.

## Introduction

The study of brain maturation and early cognitive development in humans has played a central role for the understanding of how the nervous system works. Humans are characterised by an extreme form of altriciality, which has recently evolved in response to mechanical and energetic constraints imposed by cranial capacity expansion and bipedal locomotion [1]. Newborn infants are helpless and highly dependent on their parents. The sensory and motor systems are poorly developed, partly because the degree of myelination is extremely low at birth [2].

At birth, learning, memory, and other cognitive abilities are already present, but in a rudimentary form. They gradually improve during infancy, childhood, and adolescence in parallel with the nervous system’s general maturation [3]. This is exemplified by development of numerical discrimination abilities. In the first week of life, infants discriminate a 0.33 ratio between the smaller and larger quantities, at six months they discriminate a 0.50 ratio, and at 10 months they discriminate a 0.67 ratio but not a 0.80 ratio (reviewed in [4]). Numerical acuity continues to increase throughout infancy and adolescence: six-year-old children can discriminate a 0.83 numerical ratio and adults a 0.90 ratio [5].

The cognitive development is much less studied in the other vertebrates. The ontogeny of perception, learning, and other cognitive functions has been examined in detail in only a few precocial species. The most studied species is undoubtedly the domestic chicken (*Gallus gallus*). Chicks are largely independent at birth, and their sensory and cognitive motor skills are fully developed. Research on newborn chicks revealed that they are born with a set of sophisticated cognitive abilities and that these capacities are in many cases not dissimilar to those exhibited by adult hens [6, 7].

Cognitive development of cold-blooded vertebrates has been studied mainly in the guppy (*Poecilia reticulata*), an extremely precocial teleost species. After a four-week pregnancy, guppies give birth to young that are miniatures of the adults. Newborn guppies swim in a coordinated fashion, perform predator inspection, discriminate social groups based on their numerosity, distinguish familiar from unfamiliar objects, and can rapidly learn to associate a specific object or a given number of items with a food reward [8–12].

Although altriciality characterises most mammals, birds, and teleosts, little is known about cognitive development in these species, and hence, there is currently little data available for comparison with human data. In zebrafish (*Danio rerio*), development time from fertilisation to hatching is among the shortest of all vertebrates, lasting between 48 and 72 hours. After hatching, larvae spend most of the time inactive, lying on their sides. Zebrafish start free swimming from 120 hpf (hours post fertilisation). However, it is only after 140 hpf that development of the digestive tract is completed, and they begin feeding. This period is defined as a non-feeding eleutheroembryo stage, and it is considered a post-hatching extension of embryonic development [13, 14]. In the period following the start of autonomous feeding, the zebrafish does not exhibit complex behaviours other than swimming and feeding. Attraction to conspecifics and complex social behaviour appears much later, when zebrafish are approximately three weeks old [15].

Simple forms of learning can be demonstrated in larval zebrafish. For example, 7–10 days post-fertilisation (dpf) larvae can be conditioned to avoid the darker side of their tank by delivering a mild electric shock [16]. It was shown that the visual system of zebrafish matured precociously, at the time of hatching. Larvae showed a startle response to a rapid change in light intensity at approximately 72 hpf, and tracking eye movements developed at approximately 80 hpf [17]. Little is known about the capacity to discriminate objects in the first week after hatching. Indirect evidence suggests that beginning from 14 dpf, larval zebrafish recognised a familiar stimulus based on its colour or shape [18]. In addition, at 14 dpf, larvae have marked preferences for certain visual patterns, suggesting that at this age they can discriminate their features [19].

The aim of this study was twofold. The objective of the first experiment was to determine whether one-week-old zebrafish could associate a food reward to a specific visual pattern and to validate an appetitive learning procedure for larval zebrafish. In four subsequent experiments, we used this procedure to determine the visual discrimination capacities of newborn zebrafish in tasks requiring discrimination between stimuli that differed in colour, shape, or spatial orientation.

## Results

### Experiment 1. Development and validation of an operant conditioning procedure

Twelve zebrafish larvae were trained in a two-compartment apparatus for five consecutive days (from 8 to 12 dpf), and food was delivered near one of the two visual patterns placed at the two far ends (Figure 1).

**Figure 1.**
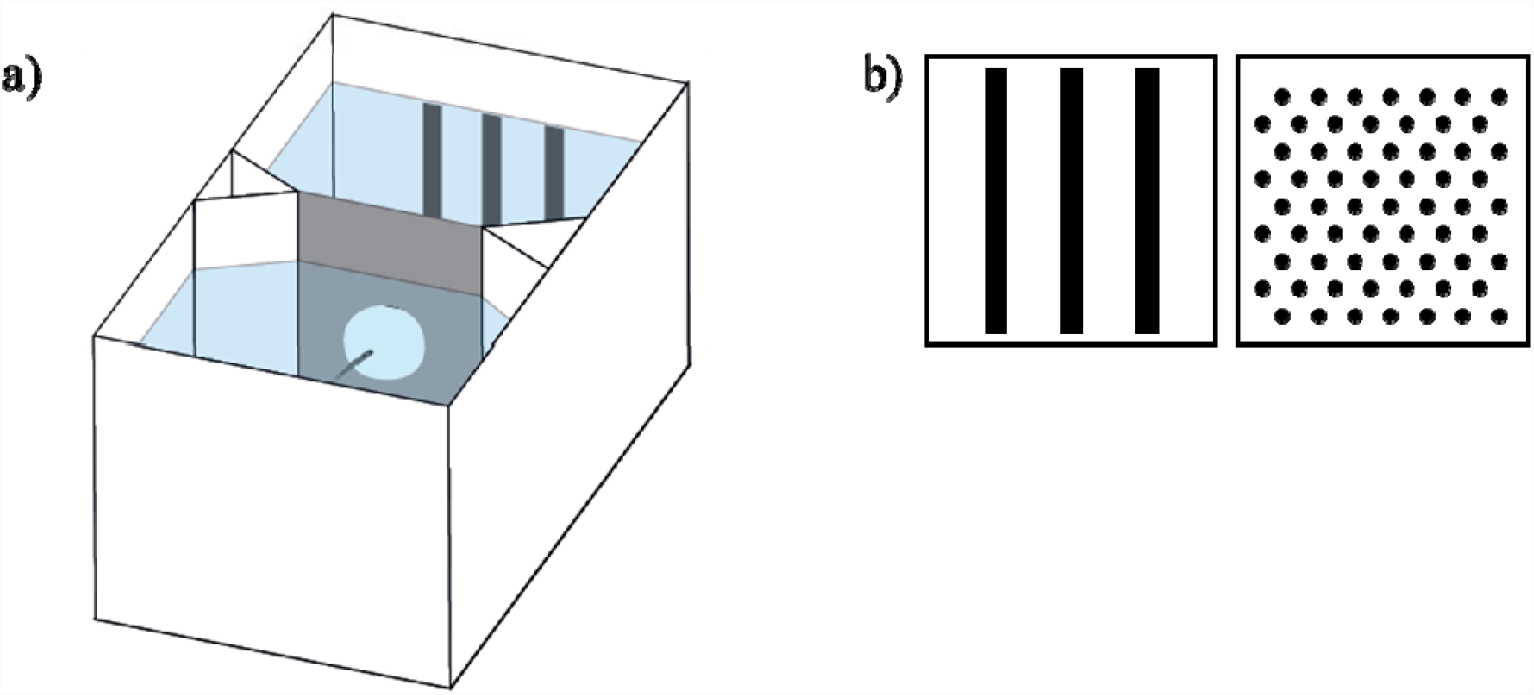
A) Apparatus used for individual training of zebrafish larvae. B) The two patterns used for the learning task of Experiment 1.

To minimise social isolation for the first two days, training was done in a group, and their behaviour was recorded. Subjects were then individually trained for the remaining three days. Performance was measured as time spent in the compartment with the positive stimulus during the 90-min period preceding food delivery. On average during each 90-min period, larvae moved 18 ± 10 (Mean ± SD) times between compartments. Subjects significantly preferred the reinforced stimulus (proportion of time 0.700 ± 0.086; one sample *t*-test, *t*(11) = 8.085, *p* < 0.001). Performance did not significantly vary across days (LMM: *F*_2,60_ = 1.662, *p* = 0.198; Figure 2), and there was no difference between larvae reinforced on vertical bars or on a dot array as positive (*F*_1,60_ = 0.002, *p* = 0.969; Figure 2). Timing of stimuli inversion had no effect (*F*_1,60_ = 0.236, *p* = 0.629), and no interaction was statistically significant (all *p*-values > 0.351). Preference for the reinforced stimulus was significant, even considering only the first day of individual training (0.625 ± 0.151; one sample *t*-test, t(11) = 2.864, *p* < 0.05).

**Figure 2.**
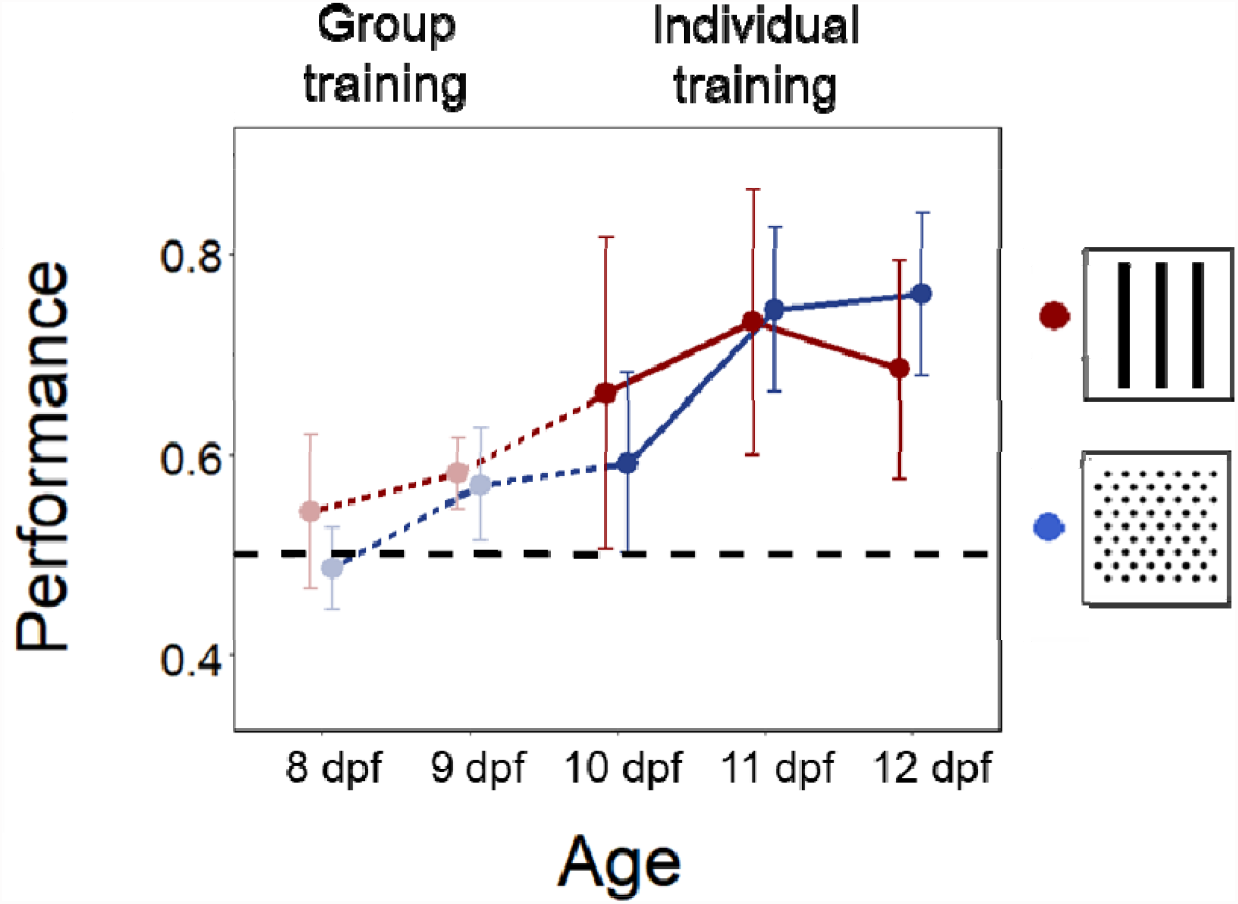
Learning performance in the five days of training (from 8 to 12 dpf). Performance was calculated as preference for the reinforced stimulus in the 90-min period preceding the delivery of the food reward. For the first two days of training (group training), preference is expressed as the proportion of larvae in the compartment with the reinforced stimulus. For the remaining three days of training (individual training), preference is expressed as the proportion of the time that a subject spends in the compartment with the reinforced stimulus. The figure shows distinct learning curves for subjects trained using vertical bars or a dot array as positive. Data points represent mean ± SEM. The black dotted line represents chance performance.

As larvae showed a significant performance already on the first day of individual training, we reconstructed the initial stages of learning by re-analyzing the videos that had been recorded in the group training phase (for the purpose of checking if larvae had learned to move between the compartments). The proportion of subjects swimming in the compartment with the reinforced stimulus was not significantly different from chance on the first day of group training (proportion in the reinforced compartment 0.514 ± 0.102, one sample *t*-test, *t*(5) = 0.328, *p* = 0.756), but it was significant on the second day of group training (0.575 ± 0.040, *t*(5) = 4.555, *p* < 0.01; Figure 2).

### Experiment 2. Colour discrimination

With the same procedure as the previous experiment, twelve zebrafish larvae were tested on a red–green discrimination (Figure 3a). The mean number of passages between compartments was 12 ± 5 (Mean ± SD). Subjects significantly preferred the reinforced stimulus (0.729 ± 0.110; one sample *t*-test, *t*(11) = 7.219, *p* < 0.001). Performance did not vary across days (LMM: *F*_2,60_ = 0.819, *p* = 0.446; Figure 3a). Larvae reinforced on the green or red did not differ in performance (*F*_1,60_ = 0.330, *p* = 0.566); timing of stimuli inversion had no effect (*F*_1,60_ = 1.265, *p* = 0.265); and no interaction was statistically significant (all *p*-values > 0.121).

**Figure 3.**
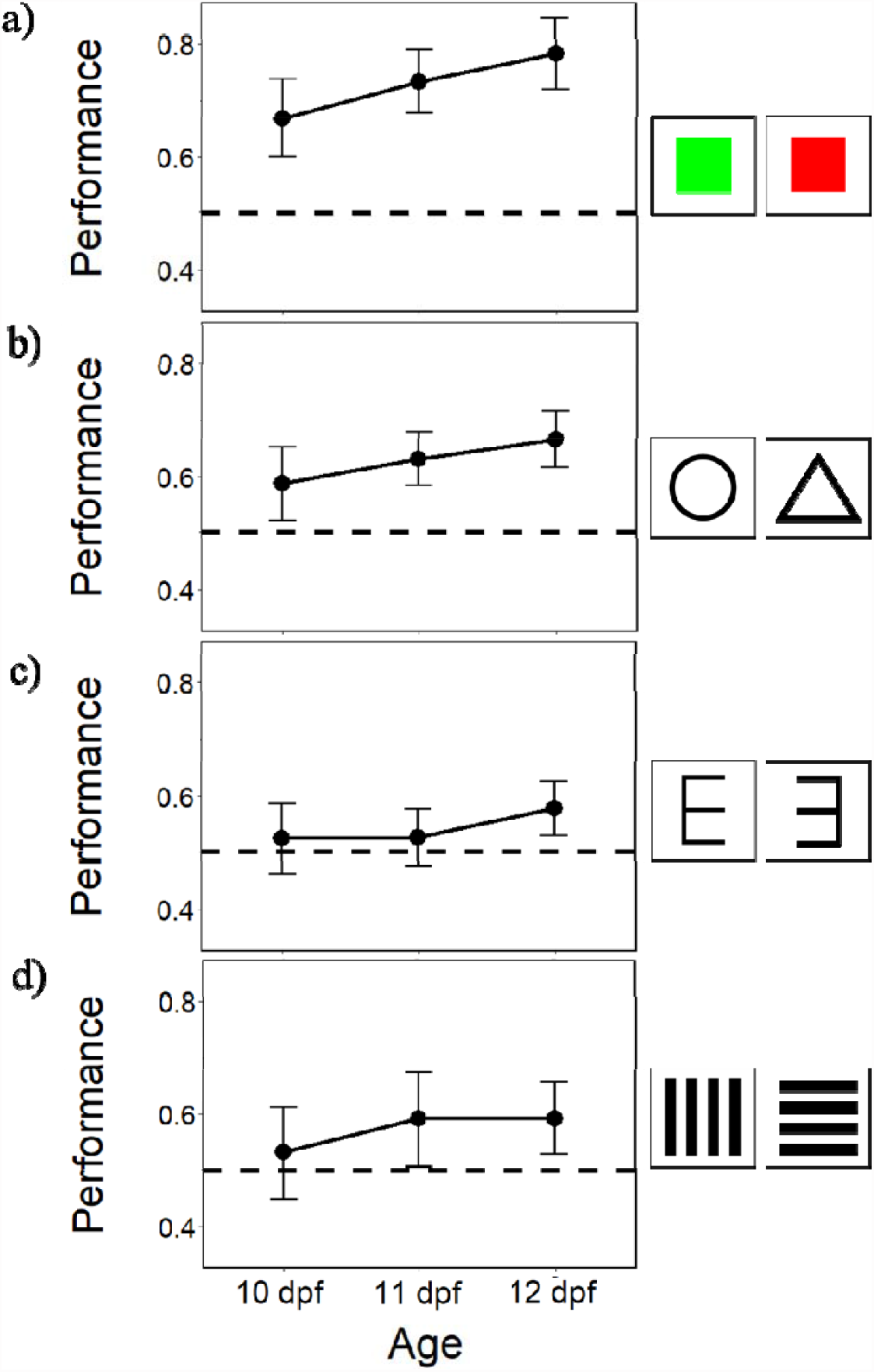
Results of Experiment 2-5. Learning curves (proportion of the time spent in the compartment with the reinforced stimulus) of zebrafish larvae in the three days of individual training. Data points represent mean ± SEM. The dotted lines represent chance performance.

### Experiment 3. Shape discrimination

Twelve zebrafish larvae were tested on a triangle–circle discrimination (Fig 3b). The mean number of passages between compartments was 15 ± 7 (Mean ± SD). Subjects significantly preferred the reinforced stimulus (0.631 ± 0.139; one sample *t*-test, *t*(11) = 3.251, *p* = 0.008). Performance did not vary across days (LMM: *F*_2,60_ = 0.571, *p* = 0.568; Figure 3b). Larvae reinforced on the triangle or circle did not differ in performance (*F*_1,60_ = 4.852, *p* = 0.062); timing of stimuli inversion had no effect (*F*_1,60_ = 0.979, *p* = 0.354); and no interaction was statistically significant (all *p*-values > 0.328).

### Experiment 4. Mirror–image discrimination

Twelve zebrafish larvae were tested in a discrimination of a figure from its mirror image (Fig 3c). The mean number of passages was 16 ± 4 (Mean ± SD). Subjects significantly preferred the reinforced stimulus (0.543 ± 0.051; one sample *t*-test, *t*(11) = 2.932, *p* = 0.014). Performance did not vary across days (LMM: *F*_2,60_ = 0.381, *p* = 0.685; Figure 3c). Larvae reinforced on E or its mirror reverse did not differ in performance (*F*_1,60_ = 0.015, *p* = 0.903); timing of stimuli inversion had no effect (*F*_1,60_ = 0.506, *p* = 0.480); and no interaction was statistically significant (all *p*-values > 0.711).

### Experiment 5. Horizontal-vertical discrimination

Twelve zebrafish larvae were tested in a discrimination of a figure from the same figure rotated by 90° (Fig 2d). The mean number of passages was 12 ± 3 (Mean ± SD). Subjects significantly preferred the reinforced stimulus (0.572 ± 0.099; one sample *t*-test, *t*(11) = 2.517, *p* = 0.029). Performance did not vary across days (LMM: *F*_2,60_ = 0.030, *p* = 0.822; Figure 3d). Larvae trained with horizontal or vertical bars had similar performances (*F*_1,60_ = 0.499, *p* = 0.483); timing of stimuli inversion had no effect (*F*_1,60_ = 0.246, *p* = 0.622); and no interaction was statistically significant (all *p*-values > 0.418).

#### Comparison of the discrimination tasks

The overall analysis on the proportion of the time spent in two choice compartments revealed that there was a significant effect of the type of discrimination test on larvae performance (LMM: *F*_3,60_ = 5.161, *p* = 0.003; Fig 4) but no effect of the day (*F*_2,262_ = 1.144, *p* = 0.320) nor of the interaction (*F*_6,262_ = 0.233, *p* = 0.965). Tukey post hoc test revealed that performance was significantly better in the colour discrimination than in the mirror-image and horizontal-vertical discriminations (both *p*-values < 0.05). No other comparison was statistically significant (all *p*-values > 0.339).

**Figure 4.**
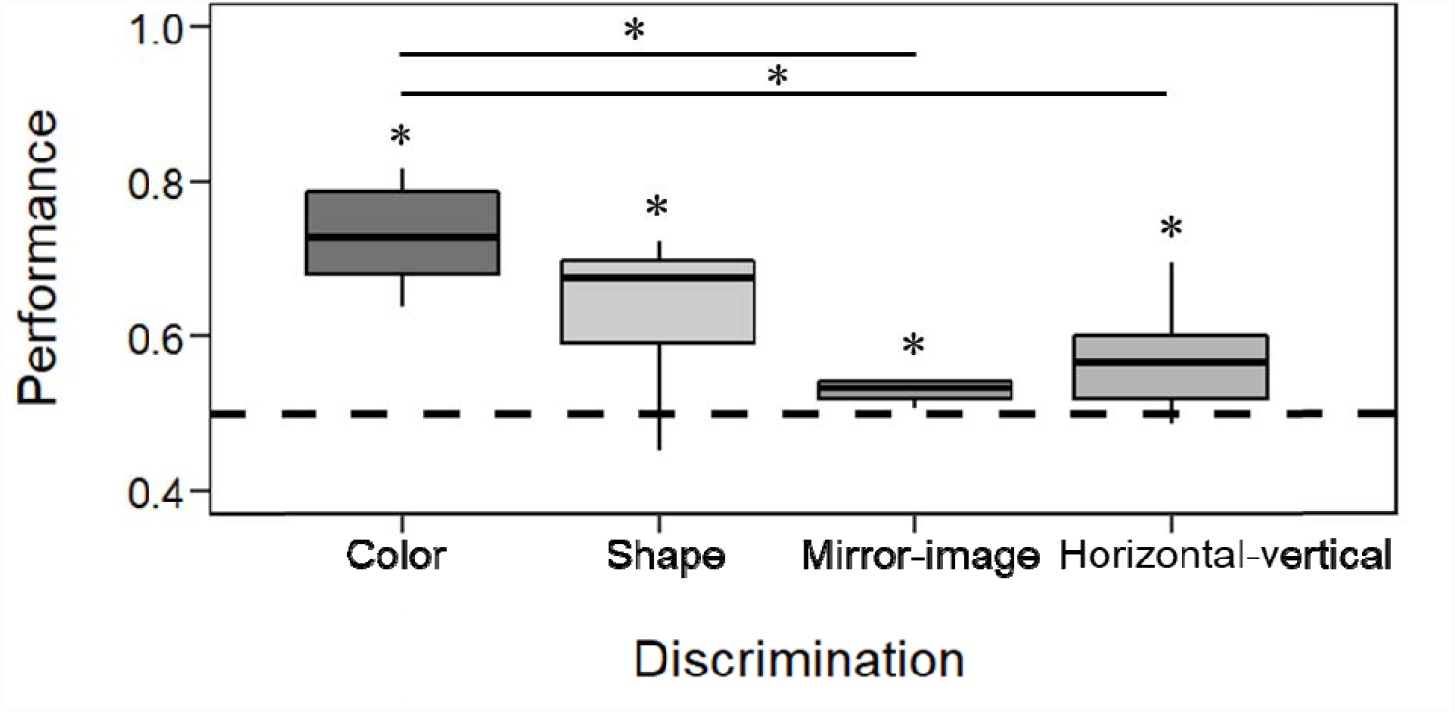
Comparison of the results of the four discrimination tasks. The box plots report median, lower and upper quartiles; whiskers represent values within 1.5 times the interquartile range. The asterisks (*) denote a significant departure from chance level (0.5).

## Discussion

Previous studies showed that newborn zebrafish can learn to respond to simple stimuli. For example, they can be conditioned to turn their tails to obtain relief from heat exposure or learn to stay away from the darker part of the tank to avoid an electric shock [16]. Using a new appetitive learning paradigm, we demonstrated that, one week after hatching, larval zebrafish rapidly associate a visual pattern with a food reward, and they can accurately discriminate two objects based on their colour or shape. Moreover, we found that young larvae can perform complex visual discrimination typical of higher vertebrates, such as discriminating a figure from its mirror image or from its 90-degree rotated version.

### Appetitive learning in larval zebrafish

The first experiment aimed to develop an operant conditioning procedure for young larvae. From 8 dpf to 12 dpf, we exposed larvae twice a day to two visual patterns, delivering food near one of the two stimuli. As consequence of this pairing, larvae increased time spent in association with the reinforced pattern. Moreover, we found no difference between subjects trained on one or the other stimulus, excluding the possibility that our results were explained by innate preferences for some patterns. To reduce the length of social isolation, the first two days of training were conducted in a group, and therefore, we have dada on individual performance only for the last three days of training. Quite unexpectedly, we observed learning to occur very early, with a significant performance observed in first day of individual training, when larvae were 10 dpf. Although we are unable to determine exactly when each subject started to learn, we re-analyzed the videos that were recorded during group training to check movements between compartments. Measuring the proportion of subjects present in the two compartments in the 90-min period preceding the delivery of food reward, we found a significant preference for the reinforced stimulus from the second day of training, at 9 dpf.

This is an especially interesting finding because, for the first time, it shows the existence of appetitive learning in zebrafish in the first week after hatching. It has been suggested that the ability of a larva to learn to avoid a dark place where it previously received an electric shock is adaptive because, in nature, this capacity allows them to avoid dangerous parts of their habitats, such as those containing predators [16]. Similarly, it is likely beneficial for recently hatched zebrafish to learn and remember a particular location or the characteristics of the microhabitat where they had previously consumed food.

The two patterns used in the first experiment differed for multiple cues (e.g., shape, size, and number of the single items composing the patterns), and larvae may have considered one of these cues or even a more basic feature of the image (e.g., the spatial frequency [20, 21]) to discriminate between them. Indirect evidence suggests that, after 14 dpf, larval zebrafish might be able to discriminate the colour and the shape of an object [18, 19]. The visual system of zebrafish starts to mature earlier, one to two days after hatching, and at 7–10 dpf, larvae have been shown to perform simple discriminations, such as recognising the dark side of their tank [17]. Given the results of the first experiment, we performed a series of discrimination learning experiments in the study’s second part to investigate perceptual abilities of zebrafish at this developmental stage.

### Colour and shape discrimination

In the five days of training, larvae learned to discriminate two figures based on either colour or shape. In both cases, the performance was very good, and there was no difference between the subjects trained on one or the other stimulus. In both experiments, we did not observe an improvement in performance during the experiment, but this may depend on the fact that we did not have information on the first two training days.

Hatching much earlier than the other teleosts, zebrafish larvae have completed only part of their development, and for the first two weeks, they exhibit a limited behavioural repertoire. However, the results of these experiments seem to indicate that some advanced cognitive functions are already present a few hours after hatching. These include the capacity to discriminate objects and memorise their characteristics and the possibility to associate specific reactions (e.g., approach or avoidance) to these objects as consequence of previous experience with them.

From this point of view, newly hatched zebrafish are not much different from newborn guppies, which are delivered after a more than 4-week gestation, and whose neural development is at a much more advanced stage [10]. The parallel with the condition of human infants is also interesting. In a context of general underdevelopment of the nervous system at birth and a complete helpless condition, some functions are more developed in human infants than would be typical for an altricial species. The senses of human infants—hearing, olfaction, and vision—are already functional at birth (and to some extent already in the fetus). Immediately after birth, human babies begin visual searching and exhibit preferences for looking at some patterns, orienting towards human faces in particular (reviewed in [22]). They show coordination of auditory and visual functions, being able to localise an auditory stimulus and turn their eyes in its direction. Newborn babies can discriminate shape colour and numerosity and show simple forms of learning, a condition that parallels what we have observed in zebrafish larvae in this study [4, 23, 24].

It is interesting to note that, in our study, only five stimulus–reward pairings were necessary to establish a significant association, and larvae reached 70–80% accuracy in colour and shape discrimination after only 10 reinforced trials. By comparison, in a recent study that used a similar appetitive paradigm and identical stimuli, adult zebrafish required five–six days, 12 trials a day, to reach similar levels of performance [25]. There may be various explanations for this apparent superiority in discrimination learning by larvae. From an ecological perspective, it is possible that, in the habitat in which larvae live, the association between food availability and particular environmental feature are stable over time, and it pays to memorise rapidly the landscape features that are associated with the likelihood of finding food. Investigations of learning and memory in insects have shown that learning speed is not necessarily related to computational capacity.

Notwithstanding their extremely small brains, species such as bees or moths can learn an association between stimulus and reward after just one or a few stimulus-reward pairings [26, 27]). Rapid learning, for example, may have evolved in bees because, once they discover a plant that produces nectar, it pays to continue to forage on the same plant species during the following hours or days [28]. Other explanations cannot be ruled out. One possible reason for faster learning in larvae is a difference in food motivation. Larval zebrafish eat a few times a day, and missing a meal may more severely affect a developing larva than an adult, which can usually starve for days without too serious consequences.

### Discrimination of stimulus orientation

In Experiment 3, we examined larval zebrafish in two discrimination tasks that are generally considered complex, the discrimination of a figure from the same figure rotated by 90° and the discrimination of a figure from its mirror image. Some species seem to have problems in discriminating rotated figures, although many others easily learn this type of task [29–31]. In adult zebrafish, the learning rate for 90°-rotated figures (the same stimuli used in the present study) matched the learning rate for colour and shape discrimination [25]. Larvae were able to discriminate between the two rotated figures, but their performances were lower than that shown in previous experiments. Therefore, the comparison with the performance of adult zebrafish seems to indicate that the ability to discriminate figures that differ in orientation is less developed at birth, and this capacity improves later in life with the maturation of the nervous system or with experience.

In the discrimination of a figure from its mirror image, the accuracy of larval zebrafish was just above chance, reaching approximately 55% preference for the reinforced stimulus by the experiment’s end, the lowest performance of the whole study. This result is not surprising, as the discrimination of mirror images was found to be an extremely difficult task for a variety of other organisms, such as octopuses, fish, rats, monkeys, and humans [32–37]. Though very low, the performance of newborn zebrafish do not appear much different from the performances observed in the adults of the investigated teleost species [25, 33, 38]. The difficulty of discriminate mirror images is commonly explained with the fundamentally bilateral symmetry of the perceiving organism’s nervous system [39, 40]. However, other authors point out that there are rare situations in nature in which it would be beneficial for an organism to discriminate mirror images, and probably, there has been little or no selective pressure for evolving this capacity [32].

## Conclusions

Rather unexpectedly for an extremely altricial species, we found that newly hatched zebrafish possess cognitive capacities that, in some domains, compare with those of the adults. A mechanism for appetitive learning adds to the other forms of associative learning previously described. Furthermore, larvae appear capable of rapidly memorising the characteristics of an object based on multiple cues and operating subtle discriminations among similar objects. Our results suggest that, as observed in human infants (reviewed in [22, 41]), in an higly altricial species some advanced cognitive functions may emerge early in brain development if these function have a high survival value at a specific developmental stage.

Zebrafish have recently emerged as an important model for understanding the mechanisms of brain functioning and for investigating human neuropathologies. Most research in this field is done on embryos and larvae before 10 dpf due to their amenability to genetic manipulation, high-resolution brain imaging, and to the possibility of high throughput *in vivo* screening. For example, mutant zebrafish lines have been obtained showing alterations of TAU protein functioning that express early in development and resemble the key pathological features of human TAU-related neurodegeneration [42]. Zebrafish models have been generated to investigate the neurodevelopmental basis of psychiatric disorders, for developing new antipsychotic drugs (e.g., [43, 44]), and for studying re-myelination process in pathologies such as multiple sclerosis or brain injuries [45]. Until now, research has been hampered by the difficulty to evidence complex behaviours and advanced cognitive functions in early developing larvae. The discovery that zebrafish can be rapidly conditioned and possess remarkable discrimination abilities within days of hatching opens the way for future developments in research, for example studying the neural organisation of complex cognitive functions or using new measures of the effectiveness of therapeutic intervention on brain diseases.

### Limitations of the Study

We describe the occurrence of appetitive learning in recently hatched zebrafish larvae. The present study, however, has several limitations. First, for ethical reasons (to reduce social isolation to a minimum), in this study, we trained larval zebrafish in a group for the first part of the experiment. The study revealed that associative learning occurred in larvae faster than we expected, and we therefore missed important information on the early stages of individual learning. Future studies should investigate this aspect by studying larvae individually from the beginning. Another limitation of our study is that we tested larval zebrafish from 8 dpf, and we did not attempt to assess the precise onset of associative learning capacities in hatchlings. This might limit our procedure’s application in neurobiological research because most brain imaging and brain mapping studies are conducted on younger larvae (usually 6 dpf), and many genetic tools have been developed for even earlier developmental stages. Future experiments would be necessary to assess our method’s validity for brain studies.

## Methods

### Experimental subjects and animal housing

The subjects were wild-type zebrafish larvae that originated from an outbred stock maintained in the zebrafish facility of the Department of Biology, University of Padova. The larvae were housed in several petri dishes (10 cm Ø, h:1.5 cm) in a solution of Fish Water 1× (0.5 mM NaH_2_PO_4_*H_2_O, 0.5 mM Na_2_HPO_4_*H_2_O, 1.5 g Instant Ocean, 1 L deionised H_2_O) and Methylene blue (0.0016 g/l). Before the experiment, they were housed in the same room at a density of approximately 30 larvae per petri dish, which were maintained at a temperature of 28.5 ± 1 °C and illuminated according to a 14:10 h light:dark cycle. Zebrafish larvae were fed twice a day with dry food (an admisture of GEMMA Micro 75 and TetraMin flakes, particle size: 0.75 mm) from the age of 6 dpf.

We tested 12 zebrafish in each experiment, for a total of 60 subjects. Throughout this manuscript, we used the standard age classification for zebrafish studies, which starts with the fertilisation day and is expressed in days post fertilisation or dpf (e.g., [46]). At the beginning of the individual test (see details below), larvae were 10 dpf, and their sex was undetermined because sexual differentiation occurs much later at approximately week 11–12 post-fertilisation [47].

### Apparatus

Two different experimental apparatuses were used for this study. The familiarisation phase and the initial group training were conducted in an hourglass-shaped apparatus (12 × 4.8 cm and 4 cm high; Figure 5; [48]), 3D printed with white PLA material, and filled with 3.5 cm of Fish Water 1×. A central corridor (4.3 cm in length) connected two identical compartments. In the middle of the corridor, a 3D printed, grey plastic panel (3 × 3.2 cm) with a central hole (1 cm Ø) allowed larvae to move from one compartment to the other one.

**Figure 5.**
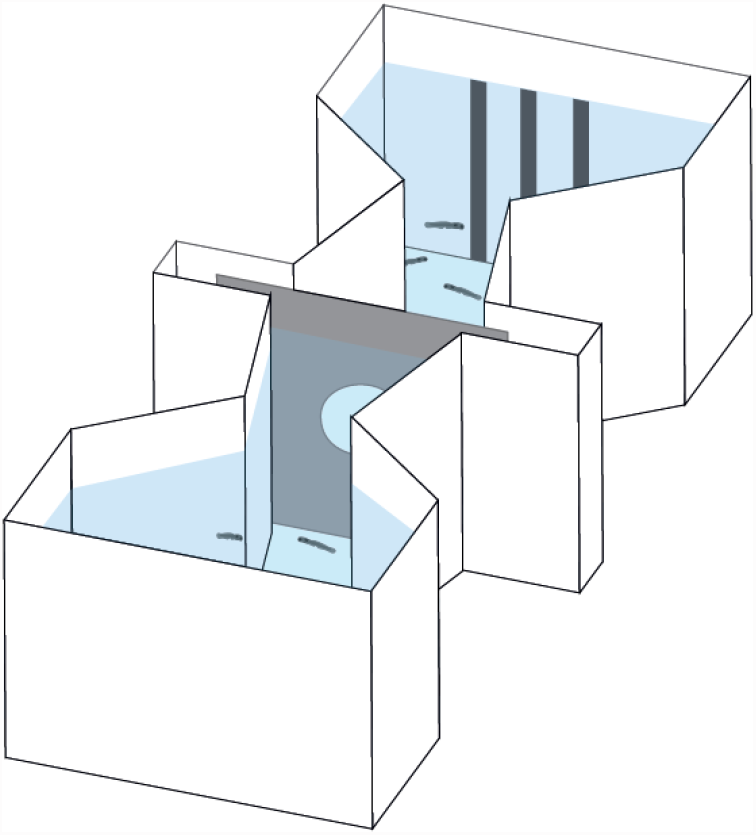
Apparatus used for the familiarization and group training phases.

The individual visual discrimination tests were conducted in a similar but smaller apparatus (7 × 4 cm and 4 cm high; Figure 1a) filled with 3.5 cm of Fish Water 1×. Four group training apparatuses were placed in a white plastic box (60 × 40 cm and 27 cm high) and lit by two 0.72 W LED lamps placed symmetrically along the major axis (Figure 6). Internally, each wall of the box was covered with white plastic sheets. An identical box was used for the individual training phase and could contain up to twelve apparatuses at the same time. Video cameras were placed above apparatuses to record larvae behaviour.

**Figure 6.**
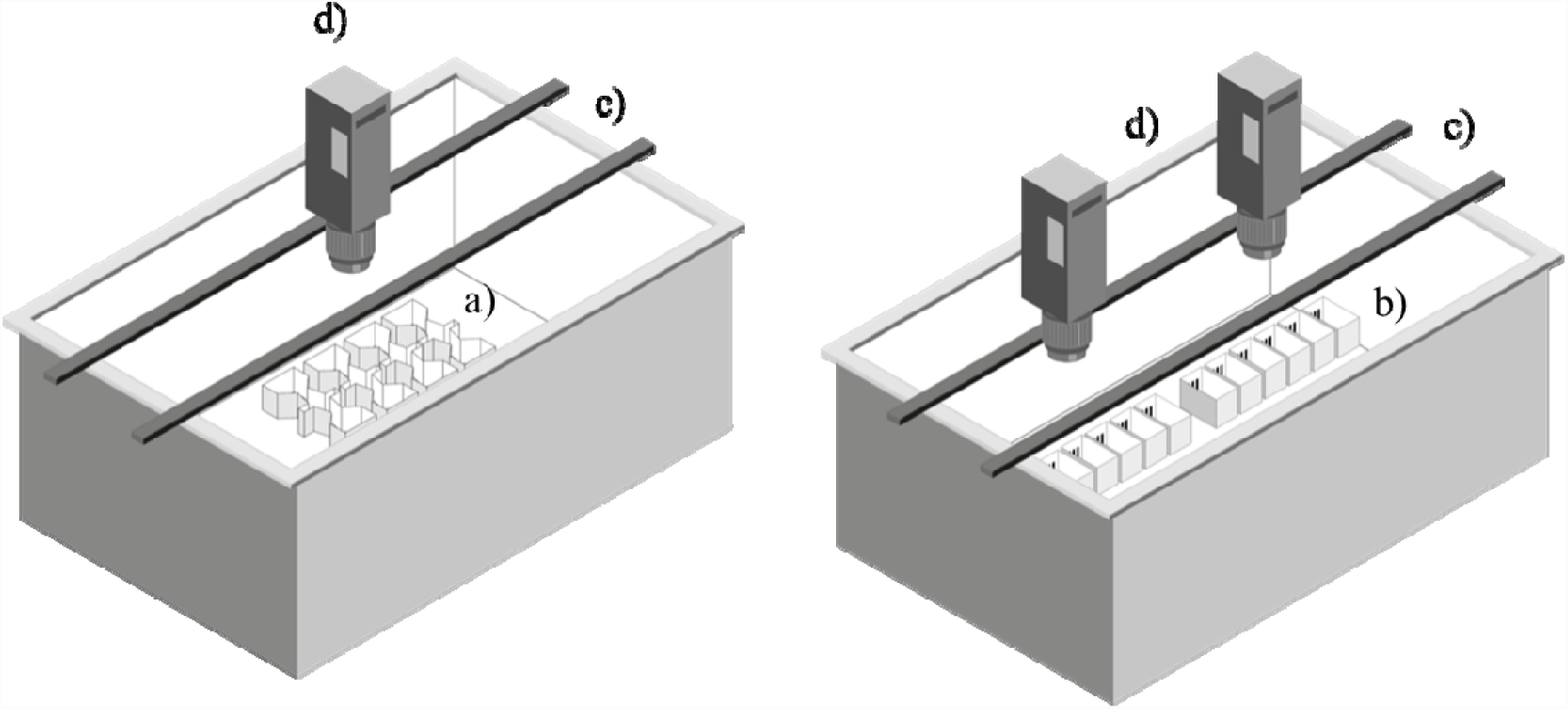
Setup for recording experiments. Four identical group training apparatuses (a) and twelve individual apparatuses (b) were accommodated in opaque plastic box to reduce any external influence. The box was lit by two lamps (c) placed symmetrically along the major axis and behaviour was recorded by videocameras (d) placed above the box.

### Stimuli

Stimuli were created using Microsoft PowerPoint and printed on 4.8 × 3.8 cm white, laminated cards. Experiment 1 aimed to develop and validate an operant conditioning procedure for larval zebrafish. To minimise the role of stimulus discrimination, we used two different visual patterns. We avoid using two hues, such as black and white, because larvae have strong preferences for the luminance of their environment. Instead, we used two patterns that differed for multiple cues (e.g., orientation and spatial frequency of the pattern, shape, size and number of the single items composing it), although they had the same percentage of black and white surfaces. One pattern consisted of three identical and parallel vertical bars (3.65 × 0.30 cm), the other consisted of an array of 63 equally spaced (0.2 cm) dots (Figure 1b).

In Experiments 2–5, we used the same stimuli used in a recent study with adult zebrafish [25]. In Experiment 2, we subjects zebrafish to a colour discrimination. The colour stimuli were one red and one green square of the same brightness (red: RGB: 255, 0, 0; green: RGB: 0, 255, 0; 2 × 2 cm; Figure 7). In Experiment 3, we tested subjects in a shape discrimination. The stimuli were one black outlined circle and one black outlined triangle (circle: Ø = 2.26 cm, area = 3.55 cm^2^, perimeter = 7.10 cm; triangle: base = 2.80 cm, height = 2.26 cm, area = 3.16 cm^2^, perimeter = 8.12 cm; Figure 7). In Experiment 4, we tested each subject in mirror-image discrimination. The stimuli consisted of the capital letter E from the Latin alphabet (height = 2.50 cm, width = 1.60 cm) and its left–right mirror-reversed image (Figure 7). In Experiment 5, we tested each subject in an orientation discrimination. The stimuli consisted of four parallel black bars (height = 2.88 cm, width = 0.4 cm) arranged vertically in one stimulus and horizontally in the other (Figure 7).

**Figure 7.**
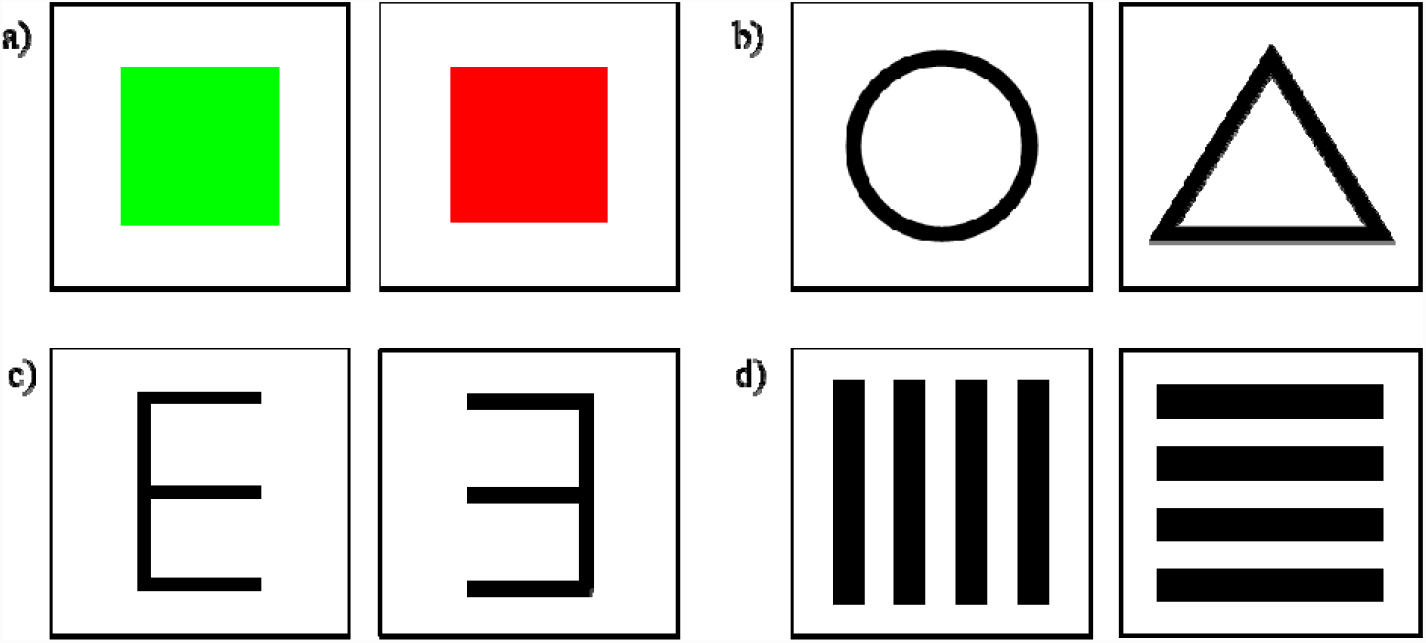
Stimuli used in Experiment 2-5.

### Procedure

The procedure was developed and validated in the first experiment and remained unchanged in the following four experiments. Previous studies suggested that social behaviour develops at approximately three weeks after fertilisation in zebrafish larvae [49]; however, it is not known whether they need conspecific visual or olfactory cues for proper maturation, as other vertebrates do, and whether they suffer from social isolation. Thus, to minimise the duration of social isolation, larvae were initially trained for two days in a group and then individually for three days. The experiment consists of three phases, an initial two-day familiarisation phase (6–7 dpf), a two-day group training phase (8–9 dpf), and a three-day individual training phase (10–12 dpf). The first two phases were conducted in the larger version of the apparatus, whereas the individual training phase was conducted in the smaller individual apparatuses. For the entire duration of the experiment, larvae were fed twice a day (h 10:30 and h 15:00).

#### Familiarization phase

Twenty 6-dpf larvae were moved using a Pasteur pipette in the familiarisation apparatus, in which no grey panel was inserted. The morning of the subsequent day, the panel with the hole was inserted to habituate larvae to pass through the hole and move between the two compartments.

#### Group training

Group training began when larvae were 8 dpf and lasted two consecutive days. Half subjects were trained on one stimulus (dots array), and the other half were trained on the other (vertical bars). Each day, we administered two reinforced trials, one in the morning and one in the afternoon (at h 10:30 and h 15:00). This was performed by leaning the two stimuli against the apparatus’s two short walls and delivering the food in the compartment with the reinforced stimulus. Stimuli remained in place during feeding and were removed 90 minutes after the delivery of the food reward. To avoid side biases, stimuli were swapped between compartments every other trial. For half of the groups, this was done before the morning trial, whereas for the other half, it was done before the afternoon trial. In experiment 1, the group training phase was video-recorded with the purpose of checking whether larvae had learned to move between the compartments. These videos were also subsequently used to measure whether larvae had started to associate with the reinforced stimulus prior to the individual training phase.

#### Individual training

At the end of the second day of the group training, larvae were moved from the group apparatus to the individual test apparatuses. The test phase lasted three consecutive days, during which we administered two reinforced trials per day with the same schedule of the group training phase and video-recorded the subjects’ behaviour for 90 minutes prior to food administration. To avoid any cueing effect due to food residuals from the previous trial, each subject was gently pipetted and moved to an identical clean apparatus at the end of each trial. The schedule for stimuli swapping was the same as the group learning phase. Overall, each subject performed 10 reinforced trials, four during the group training and six during the individual training.

### Video scoring and statistical analyses

From the video recordings of the individual test phase, we scored the time, expressed in seconds, that each larva spent in the compartment with the reinforced stimulus and the time it spent in the compartment with the non-reinforced stimulus. We also scored the number of passages between the two compartments for each subject. One-third of the videos from all experiments were scored by two different experimenters. Interrater reliability was calculated with the Spearman rank correlation and was found to be high (*ρ* = 0.968, *p* < 0.001).

Analyses were performed in R version 3.5.2 (The R Foundation for Statistical Computing, Vienna, Austria, https://www.r-project.org). We analysed the proportion of time spent in the compartment with the reinforced stimulus using one-sample *t*-tests to evaluate whether it differed compared to chance level (0.50) for each experiment. Additionally, we performed a linear mixed-effects model (LMM, ‘lmer’ function of the ‘lme4’ R package) for each experiment to assess whether the performance differed between days, between the type of reinforced stimulus, and between the type of inversion of the reinforced stimulus (between daily sessions or between days). Thus, the LMM was fitted with the serial number of the day, the type of reinforced stimulus, and the inversion as fixed factors and the individual ID as random factors. Lastly, we performed a LMM to investigate whether the larvae performance differed between the four different discrimination tests (i.e., colour, shape, mirror-image, and horizontal-vertical). This model was fitted with the discrimination and the serial number of the day as fixed effects, and the individual ID nested within the discrimination test as random effect. In case of a factor’s significant effect, pairwise comparisons were performed with Tukey post hoc tests.

For experiment 1, we also measured the number of larvae in the two compartments during the group training phase. This was done by pausing the videos at one-minute intervals and counting the number of larvae in each compartment. We checked whether the proportion of larvae in the compartments with the reinforced stimulus differed from the chance peformance level (0.50) using one-sample *t*-tests.

